# Computational identification of small molecules for increased gene expression by synthetic circuits in mammalian cells

**DOI:** 10.1101/2024.09.05.611507

**Authors:** M Pisani, F Calandra, A Rinaldi, F Cella, F Tedeschi, I Boffa, N Brunetti-Pierri, A Carissimo, F Napolitano, V Siciliano

**Author notes:** Department of Biosystems Science and Engineering (D-BSSE), ETH Zürich, Mattenstrasse 26, Basel 4058, Switzerland.

## Abstract

Engineering mammalian cells with synthetic circuits is leading the charge in next generation biotherapeutics and industrial biotech innovation. However, applications often depend on the cells’ productive capacity, which is limited by the finite cellular resources available. We have previously shown that cells engineered with incoherent feedforward loops (iFFL-cells) operate at higher capacity than those engineered with the open loop (OL). Here, we performed RNA-sequencing on cells expressing the iFFL and utilized DECCODE, an unbiased computational method, to match our data with thousands of drug-induced transcriptional profiles. DECCODE identified compounds that consistently enhance expression of both transiently and stably expressed genetic payloads across various experimental scenarios and cell lines, while also reducing external perturbations on integrated genes. Further, we show that drug treatment enhances the rate of AAV and lentivirus transduction, facilitating the prototyping of genetic devices for gene and cell therapies. Altogether, despite limiting intracellular resources is a pervasive, and strongly cell-dependent problem, we provide a versatile tool for a wide range of biomedical and industrial applications that demand enhanced productivity from engineered cells.

## Introduction

Mammalian cell engineering with synthetic circuits is a disruptive technology for the study of biological processes such as cancer treatment^1–3^ cell differentiation^4^, gene pathway elucidation^5^ and evolution^6^. Over the years it has also gained clinical relevance for the production and test of biologics^7–9^, such as recombinant proteins^10^, viral vectors, virus-like particles and cell-based therapies. At the core of these applications is the transient or stable expression of genetic payloads to facilitate rapid and robust genetic prototyping of synthetic circuits, and regulatory compliance in biomanufacturing.

We have recently shown that the dependence of gene expression on resources availability is detrimental to the operational capacity of the host cells^11–14^ and therefore to therapeutics bioproduction and genetic circuits characterization. This issue was overlooked in the first wave of mammalian cell engineering but it is now being addressed with complementary approaches that span from the characterization of circuit-host cell interaction of different genetic modules (e.g., promoters, Kozak, PolyA)^14^, which strongly depend on the cell target and may vary across cell types, to the implementation of biomolecular incoherent feed forward loops (iFFLs) controllers based on endoribonucleases or microRNAs (miRNAs)^12,13,15^ that conversely are portable and re-adaptable. Importantly, we found that in miRNA-iFFL the protein production rate increases by a mechanism of translational resource re-distribution^15^ enhancing the operational capacity of the engineered cells. However, according to the design they may require the implementation of more genetic modules and thus additional cellular resources.

On the other end, small molecules employed *ex-vivo* induce a rapid and reversible response in treated cells^16,17^. Small molecules are widely used for several biological purposes, such as fine-tune protein expression^18,19^, stem cell differentiation^20,21^, immune cells expansion for bio-manufacturing of immunotherapies^17^. The identification of small molecules typically requires the screening of thousands of compounds, which is laborious and time consuming^22^. Recently, a computational tool named “*DECCODE*” (Drug Enhanced Cell COnversion using Differential Expression) was introduced to identify small molecules that were able to regulate cellular processes towards a targeted whole-genome transcriptomic signature. DECCODE compares the expression profile of a target cellular state to those induced by a large collection of drug treatments, bypassing high-throughput screening for drug discovery. For each comparison, the method provides a similarity score measuring how much a given drug induces transcriptional features mimicking those observed in the target cells. Therefore, the approach is entirely data-driven and unbiased towards specific biological mechanism^23^.

Here, we sought to define the transcriptomic profile of cells expressing miRNA-iFFLs to be used as a target for the DECCODE algorithm, reasoning that the resulting molecules could provide an efficient and genetically non-invasive tool, to increase transgene expression rate of engineered cells. We performed RNA-sequencing analysis to characterize the transcriptional signature of cells expressing a two-genes system either in the more burdensome open loop architecture (OL-no miRNA regulation) or in the miRNA-iFFLs architecture. By comparing thousands of transcriptomes from drug-treated cells with those generated by OL and miRNA-IFFLs designs, DECODDE selected drugs with profiles similar to those of miRNA-iFFLs and divergent from the OL (**Fig.1**).

**Figure 1.**
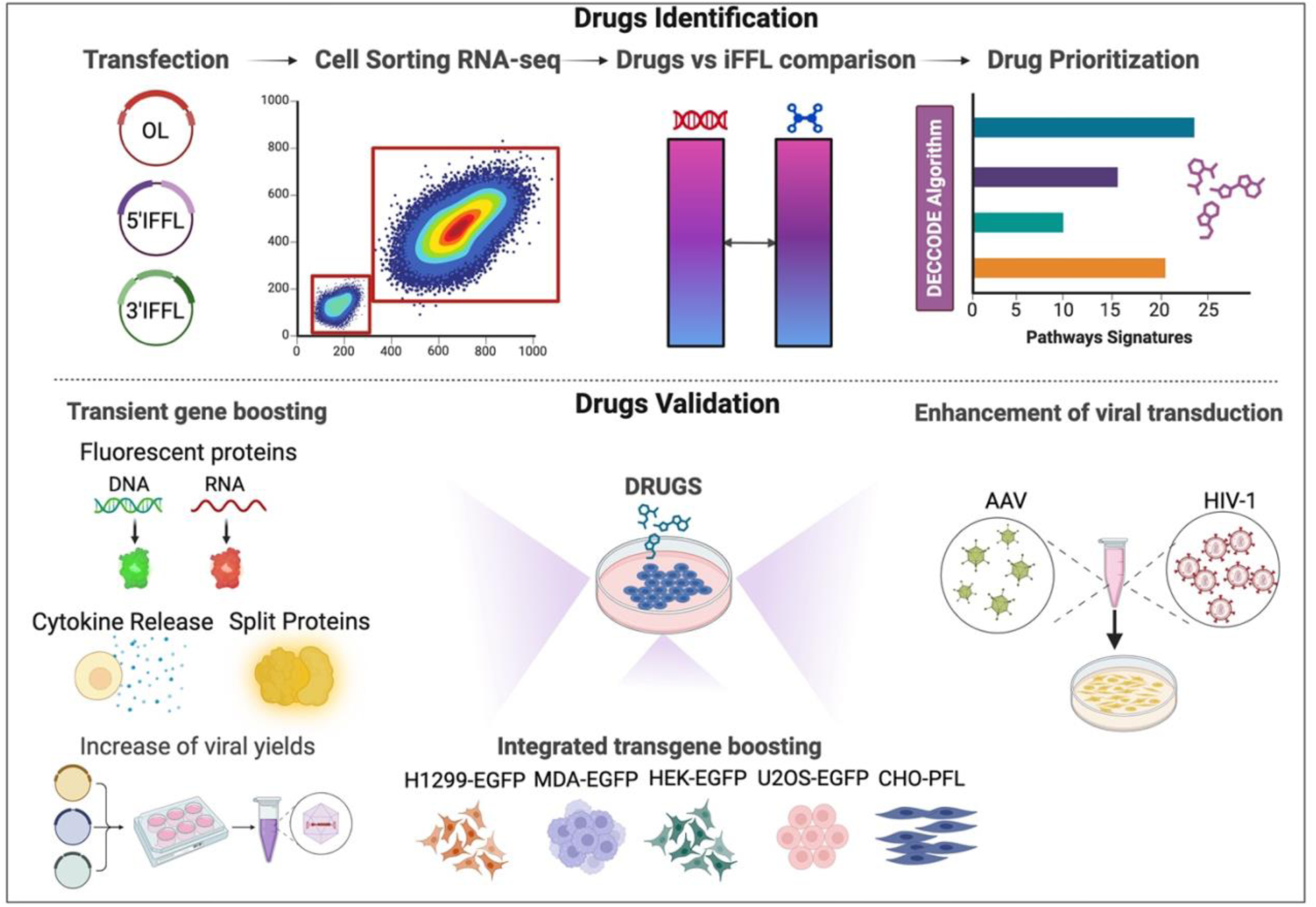
Pipeline of the study. **Top**: drug identification. H1299 cells transfected with EGFP and mKate fluorescent proteins in different circuit topologies, Open Loop (OL) and microRNA-Incoherent Feed-Forward Loop (miRNA-IFFL) were sorted according to fluorescence expression and RNA was extracted to carry out RNA-sequencing. A gene signature matrix was thus generated and converted into pathway signatures. Drug prioritization is obtained by DECODDE algorithm, comparing the transcriptomes of thousands of drug-treated cells with those generated by OL and miRNA-IFFL designs, to select drugs with profiles similar to those of miRNA-IFFLs. **Bottom**: drugs validation. Increased protein expression was evaluated and exploited in different experimental settings, including expression of genetic constructs encoded by DNA or RNA in transient (left), and integrated cell lines (middle). Finally, on a more applicative exploitation of DECCODE,we evaluated the effect of the leading drugs on viral transduction. *Created with BioRender.com*.

We tested the top compounds in several experimental settings and various cell lines including transient transfection of DNA or RNA or stable gene expression (**Fig.1**). Finally, we demonstrate that drug treatment enhanced *in vitro* adeno-associated virus (AAV) and lentivirus transduction, tackling current limitation of viral-delivered gene circuits screening *ex vivo*. Collectively, we show that by selecting small molecules in a black-box fashion, with no prior information or investigation about the mode of action, we have a versatile tool for a variety of biomedical and industrial applications that requires higher productive capability of the engineered cells.

## Results

### Identification of small molecules to increase the expression of exogenous genetic payloads by DECCODE

We have previously shown that exogenous DNA impose a burden on the host cells due to the finite amount of transcriptional and translational resources, resulting in a limiting productivity capacity^12,13^. We thus designed context-aware genetic networks in which miRNAs were successfully exploited as burden mitigators^12^, and observed upregulation of protein expression in the targeted cells. To better understand the transcriptional changes occurring with the different design frameworks, we co-transfected H1299 cells, the testbed of our previous studies^12–15^, with a bi-directional plasmid encoding EGFP and mKate, either in an open loop circuit architecture (OL) that lack miRNA regulation, or in the form of a miRNA-iFFL, placing target sites for miR31, highly expressed in this cell line, in the 3’ (iFFL-3’) or 5’(iFFL-5’) UTR of mKate (**Fig.2a**). We next sorted the cells that were either non-transfected (NTr) or transfected (Tr) according to fluorescence expression, to compare the OL that are more sensitive to burden, with iFFL-3’ and iFFL-5’ architectures.

We performed Principal Component Analysis (PCA) to investigate the distribution of NTr and Tr populations in a two-dimensional Cartesian space for the three transfection conditions (**Supplementary** Figure 1). In the OL design, NTr and Tr populations are spatially separated (**Supplementary** Figure 1a), while in the iFFL-3’ and iFFL-5’ configurations, the distance between populations progressively decreases (**Supplementary** Figure 1b-c). We next carried out the differential expression analysis on the three experimental groups (**Fig.2b**). As compared to iFFL samples, the OL exhibits the highest number of differentially expressed genes (**Fig.2b, red circles**), indicating the largest perturbation to the cells (**Supplementary Table 1**). This finding is consistent with the observation that iFFL topology mitigates gene expression burden and, in agreement with our previous reports, when miR-31 target sites are located at the 5’UTR the system is more resistant to genetic perturbations, showing further reduced differential expression (**Fig.2b, purple circles**) as compared to the 3’UTR (**Fig.2b, green circles**). Next, we analyzed the biological pathways enriched based on the differentially expressed genes (FDR ≤ 0.1) across the different transfection conditions. As expected, the number of dysregulated pathways is greater in OL samples (**Supplementary Table 2**), and several involve RNA processes, translation and post translational modification that are bottlenecks in protein production, indicating a correlation with resource competition and burden (**Fig.2c**). Further, many biological processes include metabolism and energy production, crucial to cell fitness and activity. Other pathways relate to innate response to external nucleotides and viral particles (RIG-I signaling pathway), suggesting a possible limitation to the expression of exogenous payloads (**Fig.2c**). Interestingly, RNA processing and translation pathways are still dysregulated in the 3’iFFL, but not in the 5’UTR, in agreement with the ribosome redistribution observed in our previous report^15^, whereas pathways associated with the innate immune response are still present in both 3’ and 5’ iFFLs. Having confirmed the different transcriptional signature in OL *versus* iFFL expressing cells, we next used DECCODE algorithm to identify small molecules that can increase gene expression of exogenous payloads. As required by the DECCODE approach, a single transcriptional model of burden mitigation was obtained by comparing cells carrying the OL or the iFFLs (**Fig. 2d**). We merged all the profiles corresponding to cells with a significant transfection level, thus diluting the main source of variation in the data. The obtained differential profile was converted to a pathway-based expression profile based on the Gene Ontology – Biological Process collection. Of note, this analysis is substantially different from the previous (**Fig.2b-c**), since it only considers the differences across the transfected conditions (OL, 3’iFFL, 5’iFFL), as compared to those intra-sample. Comparison against ∼19,000 drug-induced pathway-based expression profiles from the Library of Integrated Network-Based Cellular Signatures (LINCS)^24^ was performed to prioritize the corresponding molecules (**Fig.2d**). Specifically, small molecules inducing a profile that was similar to the miRNA-iFFL mitigated profile were ranked higher (**Fig.2e** visualizes the top 30 hits based on pairwise structural distance). We performed a Drug-set Enrichment Analysis (DSEA)^25^ of the top 30 ranked drugs to identify the common pathways on which those drugs have an impact and then compared them with the biological processes enriched in the RNA-seq data. Interestingly, several pathways related to RNA processes, protein translation and metabolism were affected by the drugs (**Fig.2f**). The DSEA of the top 30 drugs suggests that the selected drugs may affect pathways involved in resource reallocation, to effectively increase gene expression in the target cells. Finally, out of the top thirty we selected sixteen drugs based on commercial availability and tested their effects on fluorescent protein production in the OL configuration, the most burdensome to the cells.

**Figure 2.**
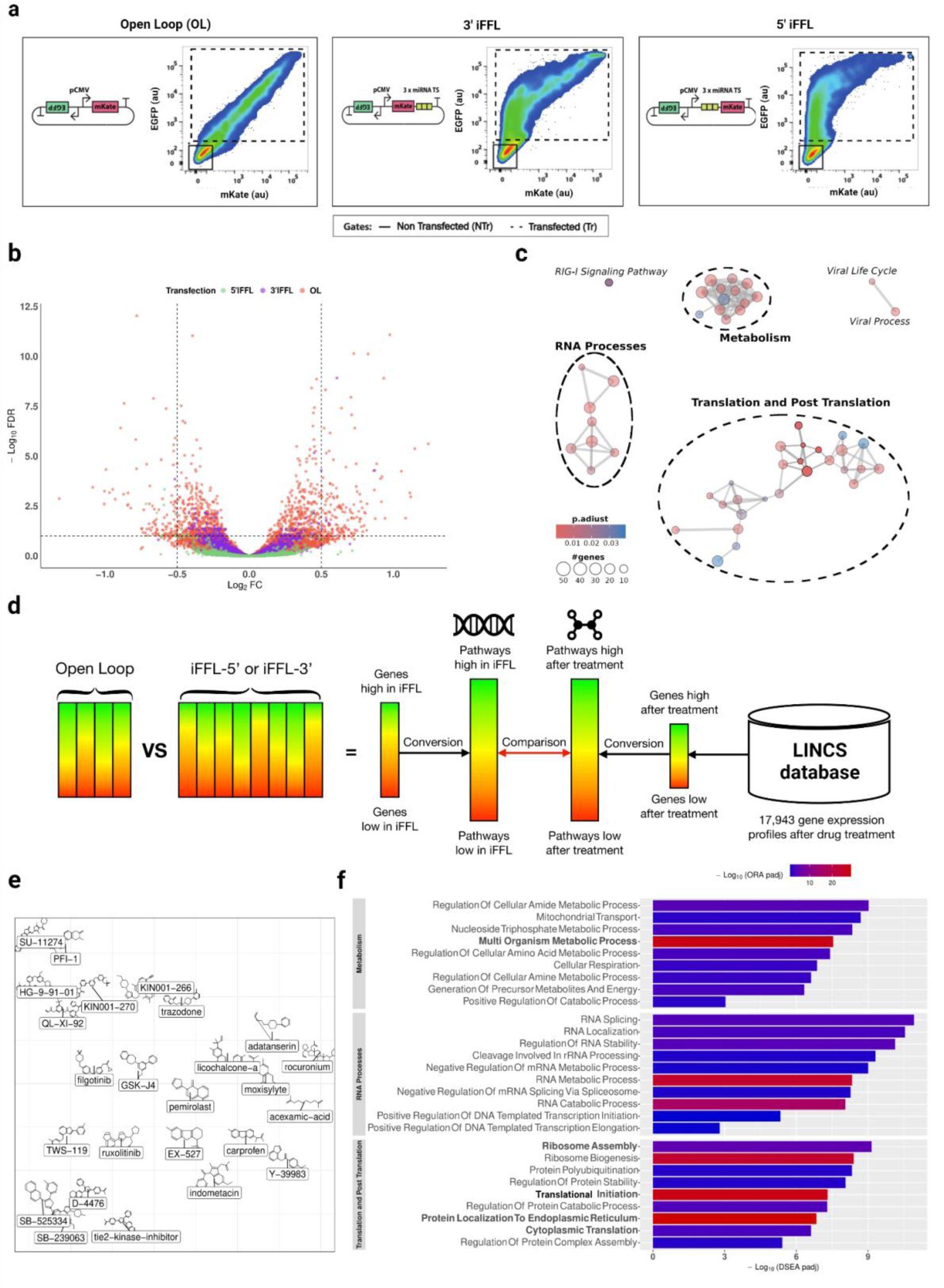
The transcriptomic landscape of burdened cells and drug identification by DECCODE. (**a**) Flow cytometry analysis of H1299 cells transfected with plasmids featuring three distinct circuit architectures: Open Loop (OL), 3’ miRNA-iFFL, and 5’ miRNA-iFFL. The 3’ and 5’ miRNA-iFFL include three target sites for miR31 which is expressed in the H1299 cell line. The dot plots illustrate the differing distributions of mKate fluorescent cell populations across the various circuit topologies. (**b**) Volcano plot depicting differential gene expression across the three circuits architecture, with the OL (red dots) showing the highest number and broadest range of DEGs. (**c**) Network visualization of significantly dysregulated pathway in cells transfected with the OL, including RNA processes, metabolism, translation and post-translation modifications. (**d**) Schematic workflow of DECCODE. The algorithm first compares differentially expressed genes in OL and iFFL transfected cells, identifying enriched pathways, which are then matched with drug-associated pathways using the LINCS database. (**e**) Visualization of the top 30 small molecules identified through DECCODE as selected by similarity to gene expression profiles observed in iFFLs. Spatial placement of the molecules is based on pairwise structural distance. (**f**) Common dysregulated pathways by Drug-set Enrichment Analysis (DSEA) of the top 30 drugs (padj as bar length) and by over-representation analysis (ORA) of cells transfected with the OL (padj as bar color); pathways also enriched in iFFL designs are highlighted in bold.

### Selected small molecules increase expression of genes delivered as DNA or RNA in different cell lines

We co-delivered EGFP and mKate encoding plasmids in H1299 cells followed by treatment with the selected compounds 4h post-transfection. The timing was chosen based on plasmid uptake that occurs relatively early after transfection (within the first few hours)^26^. Four of the sixteen drugs, namely Ruxolitinib^27^, TWS119^28^, Filgotinib^29^ and Tie2 kinase inhibitor 1 (TIE2)^30^, increase fluorescence expression by ∼10-50% compared to untreated cells (**Fig. 3a, Supplementary** Fig.2). Ruxolitinib and TWS119 had the most significant effect, while Filgotinib slightly increased mKate (∼10%) but not EGFP expression which seemed slightly reduced.

In a recent work, we optimized the simultaneous gene expression of EGFP and mKate by combining various genetic modules (promoter, kozak, polyA) in the plasmid designs^14^. This optimization was particularly effective when combining the HGHpolyA (EGFP-HGHpA) with the SV40polyA (mKate-SV40pA) in a 1:1 ratio in HEK293T and CHO-K1. We confirmed that this genetic combination improves the expression of both fluorescent proteins also in H1299 (**Supplementary** Figure 3a) and, when the four small molecules were added, data showed a further increase in the expression of both EGFP-HGHpA and mKate-SV40pA (**Supplementary** Figure 3b). In the case of EGFP-HGHpA encoding plasmid, Filgotinib treatment resulted in higher protein production than untreated cells, in contrast to what observed with EGFP bearing the SV40polyA (**Supplementary Fig.3b**) suggesting that combining genetic modules with drug treatment may enhance optimal gene expression. Furthermore, by steadily increasing the EGFP-HGHpA molar ratio, we found that this genetic design combination was also the least perturbing mKate expression, ensuring sustained levels of both genes across the different molar ratio, as compared to the EGFP-SV40polyA design (**Supplementary** Fig.4). Another important consideration is that the burden imposed by exogenous payloads on intracellular resources and genetic design is highly cell-context dependent. Therefore, we tested the effect of drugs treatment in HEK293T and CHO-K1 cells, which are the workhorses of *in vitro* biological studies and bioproduction. Our data show that protein levels of both EGFP and mKate increase by ∼30-60% in HEK293T cells (**Fig.3b**), whereas no significant differences were observed in CHO-K1 cells (**Supplementary Fig.5a**). Filgotinib, which had a mild effect in H1299 cells, emerged as the leading candidate in HEK293T cells, whereas Ruxolitinib increased protein production in both cell lines.

**Figure 3.**
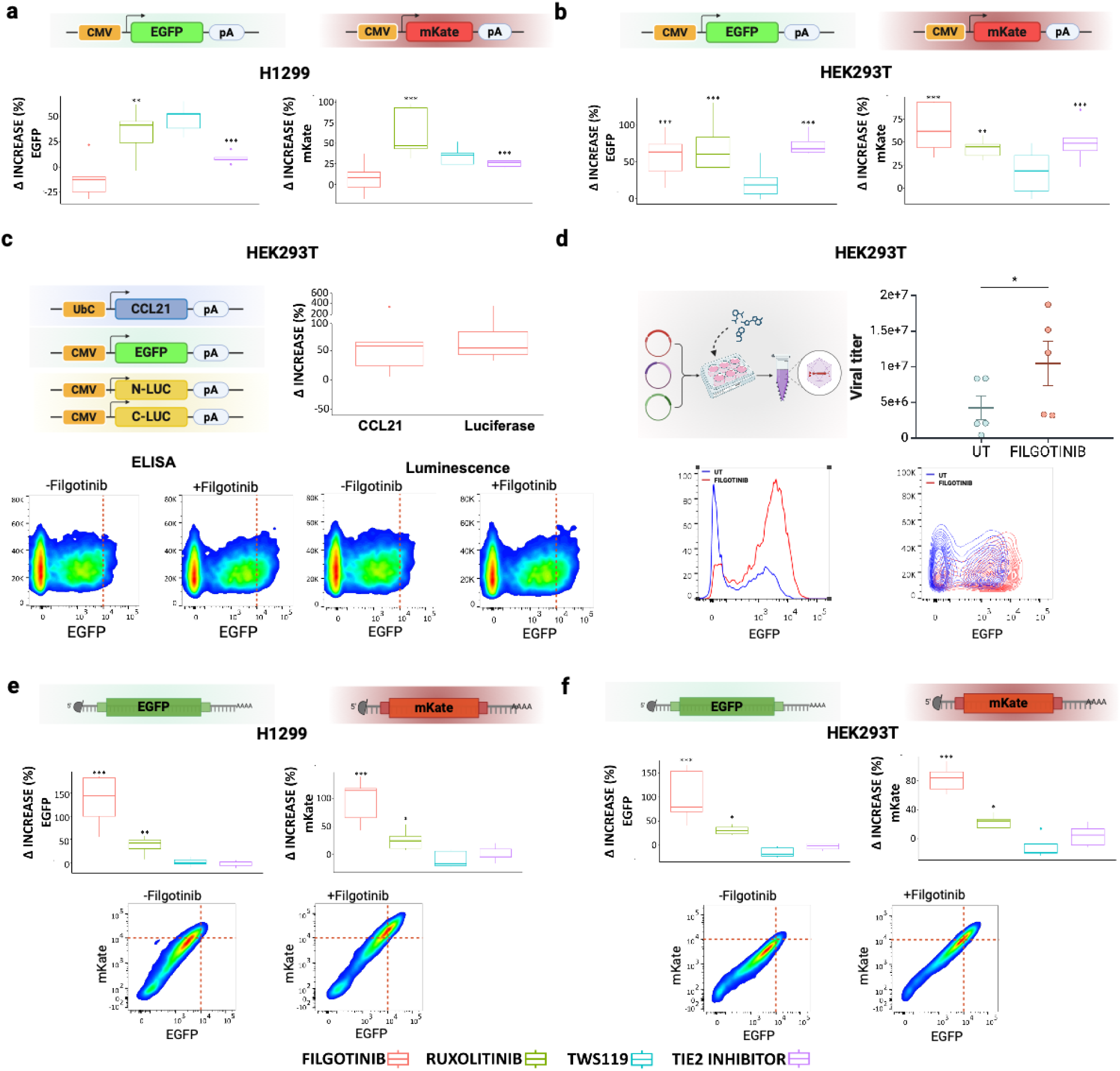
Effect of drug treatments on transient gene expression. (**a-b**) H1299 and HEK293T cells were transfected with plasmids encoding EGFP and mKate and then treated with the compounds. The boxplots illustrate the percentage increase of fluorescence intensity (drug color-code at the bottom of the panel). (**c**) Filgotinib treatment increased the release of the chemokine CCL21 and, in a split luciferase system, enhances Luciferase reconstitution and subsequent activity in HEK293T cells. The boxplot shows the percentage increase in CCL21 expression and luminescence following drug treatment. The density plots display EGFP fluorescence (co-expressed with CCL21 or with split-Luc) with and without Filgotinib treatment. (**d**) Filgotinib treatment during lentiviral production process significantly increases the lentiviral titer. The histogram and density plots illustrate the differences in number of transduced cells and fluorescence intensity w/wo Filgotinib. (**e-f**) RNA-encoded EGFP and mKate expression increases upon drug treatment, in particular with Filgotinib in H1299 and HEK293T cells. Transient transfection was performed with modRNAs. The density plots below illustrate the clear shift in fluorescence intensity with and without Filgotinib treatment. ANOVA test for statistical analysis (*p<0.05,**p<0.01,***p<0.001); N=5 biological replicates. *Created with BioRender.com*.

We then investigated whether the increased gene expression with Filgotinib and Ruxolitinib would translate into more functional outcomes, such as improved protein secretion, and protein reconstitution by an AND gate-like circuit. These experiments were performed in HEK293T cells, which are commonly used for genetic circuit prototyping, and cell engineering. For protein secretion we opted for chemokine C-C motif ligand 21 (CCL21)^31^, a cytokine that is involved in immunoregulatory and inflammation processes that we had available in the lab. CCL21 chemokine production was evaluated by ELISA assay, that showed an increase by 50% upon Filgotinib treatment (**Fig.3c**). We next co-expressed a luciferase-encoding gene split on two separate plasmids and the reconstitution was also enhanced by 50% with the same small molecule (**Fig.3c**). In both experiments we co-transfected an EGFP encoding plasmid as reference (**Fig.3c bottom**). On the contrary, Ruxolitinib did not show improvements in these settings despite increasing EGFP expression (**Supplementary** Fig.6). Since Filgotinib showed the most consistent effect in HEK293T cells, we sought to investigate whether lentivirus production, widely used to engineer different cell lines could benefit from the treatment. Lentiviral vectors are produced by transient expression of transfer and accessory plasmids in HEK293T cells. However, even if they can embed quite large genetic payloads, the efficiency of the production is inversely proportional to the size of the DNA fragments, a limitation for the engineering of genetic designs with multiple transcriptional units.

We co-transfected HEK293T cells with a lentiviral vector expressing EGFP and accessory plasmids following well established protocols^32^. We then added the drugs 4h post-transfection and collected the virus 72h later, setting a control sample without drugs. Upon virus collection, we transduced HEK293T cells, and calculated the titer based on the population of EGFP-positive cells (see Material and Methods). The results show that the treatment with Filgotinib increased the vector yield of 60% as compared to the untreated ones (UT) (**Fig.3d**).

Lastly, as RNA-encoded circuits have emerged as a novel gene regulatory modality that addresses limitations of insertional mutagenesis and immunogenicity^33,34^, we investigated whether drug treatment would benefit their expression. We co-transfected modified RNAs (modRNA) expressing EGFP and mKate and measured protein expression via flow cytometry 24h post-delivery. Both H1299 and HEK293T cells remarkably increase EGFP and mKate expression by 100% with Filgotinib (**Fig.3e-f**). Interestingly, CHO-K1 cells also exhibited higher protein expression from RNA-encoded circuits when treated with Filgotinib and Ruxolitinib (**Supplementary Fig.5b**), suggesting that these drugs may influence post-transcriptional processes. Overall, our findings indicate that Filgotinib and Ruxolitinib significantly boost gene expression and protein production across various experimental setups by DNA- or RNA-encoded genetic circuits.

### Impact of small molecules on cell engineered by stable integration of transgenes

While transient transfections are typically performed for rapid prototyping of genetic circuits, stable integrations are used in industrial and biomedical assets. Therefore, we investigated the response to drugs of stably integrated genes either in steady conditions (**Fig. 4a top right**), or when undergoing perturbations by exogenous payloads provided on-demand (**Fig.4a bottom right**).

**Figure 4.**
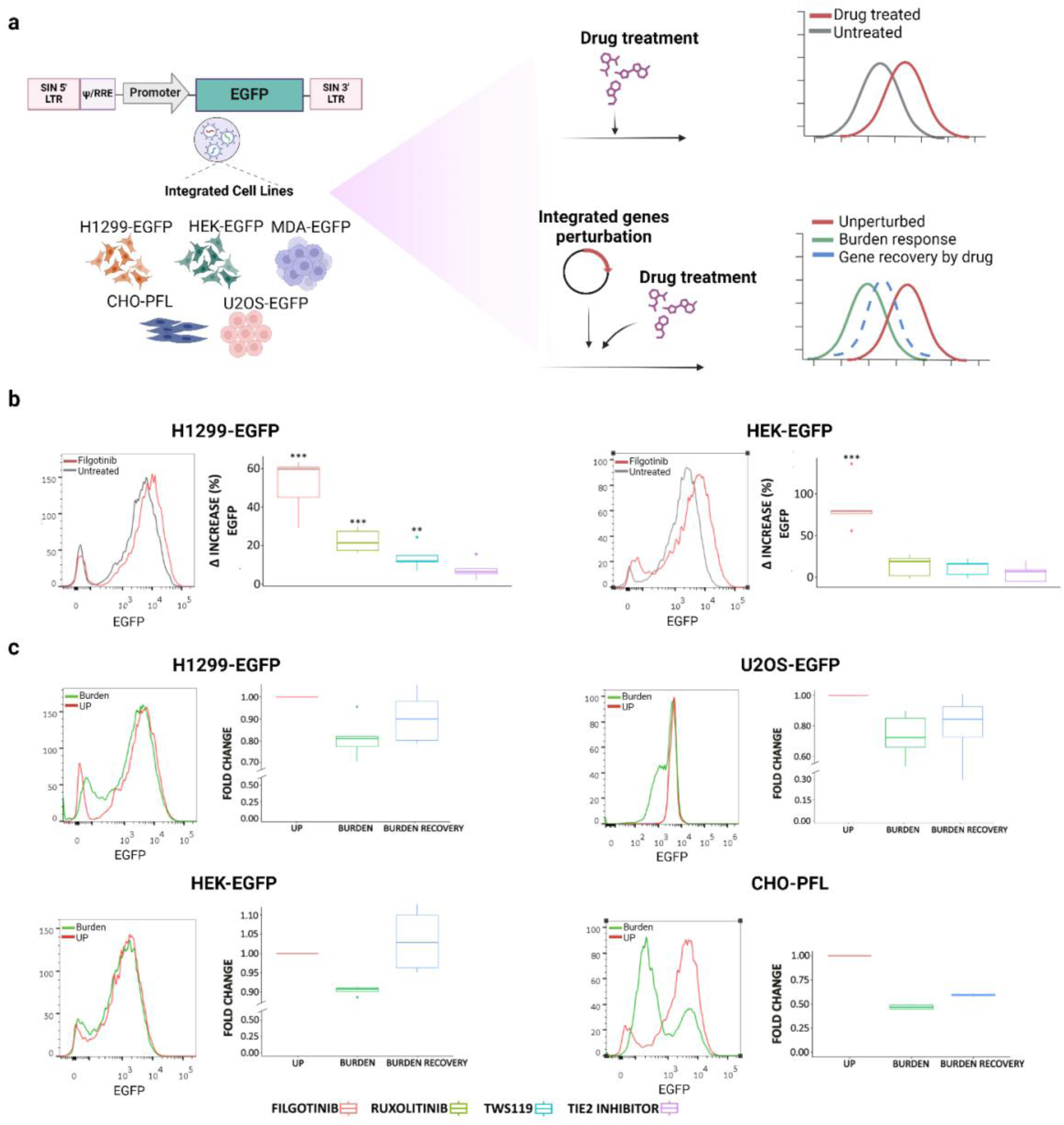
Effects of drugs treatment and transient perturbations on stably integrated genes in different cell lines. (**a**) Schematic representation of the experimental design. Cell lines H1299-EGFP, HEK-EGFP, MDA-EGFP, CHO-PFL, and U2OS-EGFP were generated by lentiviral transduction of a constitutive promoter driving EGFP. (**top right**) Cells were then treated with the compounds and changes in EGFP expression were analyzed by flow cytometry. (**bottom right**) Burden on integrated cells was applied by perturbation with a transiently transfected gene, and the response on EGFP expression and its recovery by the treatment with the drugs was evaluated by flow cytometry. (**b**) The four drugs increased expression of integrated payloads in H1299-EGFP and HEK-EGFP cell lines, in particular with Filgotinib. The histograms on the left compare EGFP expression levels in Filgotinib-treated versus untreated cells. The boxplots illustrate the percentage increase in EGFP expression following drug treatment. (**c**) Analysis of EGFP expression variations in H1299-EGFP, U2OS-EGFP, HEK-EGFP, and CHO-PFL cells under burden (green boxplot) and subsequent recovery by Filgotinib treatment (blue boxplot). Histograms depict EGFP expression profiles for cells unperturbed (UP) and on burden condition. Box plots illustrate the fold change in EGFP expression, highlighting the recovery effects of the various drugs. ANOVA test for statistical analysis (*p<0.05,**p<0.01,***p<0.001); (**b**) N=5 biological replicates. (**c**) N=4 biological replicates. *Created with BioRender.com*.

We first engineered H1299, HEK293T, U2OS, and MDA-MB231 cell lines with a lentiviral vector encoding a UbC promoter driving the EGFP (**Fig. 4a, Supplementary** Fig.7). We assessed the impact of drug treatment on integrated gene expression by adding the four compounds to cultured cells, followed by flow cytometry analysis. H1299-EGFP and HEK-EGFP cells increased EGFP expression by 10-70%, with the highest upregulation induced by Filgotinib (**Fig. 4b**), whereas U2OS-EGFP, and MDA-MB231-EGFP did not show any response (**Supplementary** Fig.7). To account for genetic payload diversity, we also developed a HEK293T cells line expressing a shEF1a driving the mCherry and demonstrated similar trend of gene expression following the treatment with Filgotinib, Ruxolitinib, TWS119 and Tie2 inhibitor (**Supplementary** Fig.8).

Since we have previously demonstrated that limiting resources can impact endogenous genes upon synthetic genes transfections^12^, we next investigated whether integrated genes are also perturbed by transient expression of exogenous payloads and if so, whether the drugs could mitigate the effect. In fact, recent work has demonstrated decreased expression of integrated genes when perturbed with plasmids that include promoters of varying strengths^35^. However, those studies focused on landing pad integrated genes, typically featuring a single locus integration, whereas we evaluated the effect upon lentiviral transduction, commonly used in various applications including cell-based therapies, which often require multiple copies of the integrated gene.

We carried out tests in H1299-EGFP, U2OS-EGFP, HEK-EGFP and MDA-MB-231-EGFP, and our results indicate that also lentiviral-transduced cell lines suffer burden and confirm cell-type dependency of this effect. Notably, H1299-EGFP, and U2OS-EGFP cells showed a decrease (between 20-45%) upon burden perturbation by CMV-mKate expression (**Fig. 4c, green boxplot)**. Conversely, HEK-EGFP exhibited only a 10% reduction (**Fig. 4c bottom, green boxplot**) and MDA-MB-231-EGFP did not vary protein expression (**Supplementary** Fig.9). In addition to cell lines engineered with a simple promoter-reporter gene, we sought to investigate the response to drug treatments and genetic perturbation of a more complex circuitry design such as the positive feedback loop, a recurrent motif in gene regulatory networks. To this end we used CHO TET-OFF cells, stably expressing the tetracycline-controlled trans-activator (tTA), engineered with a CMV-TET promoter driving the expression of tTA fused to the d2EYFP^36,37^ (CHO-PFL). In a previous report we extensively characterized the features of such motifs, as compared to the counterpart lacking the PFL^36^. CHO-PFL exhibit low d2EYFP upregulation to Filgotinib and Ruxolitinib (**Supplementary** Fig.10), and are strongly perturbed by burden (**Fig.4c, green boxplot**).

Finally, we tested whether Filgotinib, that induced the strongest upregulation of integrated genes, was also able to reduce the burden by exogenous perturbation. In H1299-EGFP, HEK-EGFP, CHO-PFL and U2OS-EGFP cells, transfection followed by Filgotinib treatment resulted reduced perturbation by CMV-mKate. In H1299-EGFP and HEK-EGFP cells, the expression approximates to baseline levels **(Fig.4c, blue boxplot)**, whereas in U2OS EGFP and CHO-PFL cells, which were less responsive to the drug, the increase was modest **(Fig.4c, blue boxplot).** These results are perhaps the consequence of enhanced protein expression by Filgotinib supplementation. This is supported by the observation that the mKate levels remain stable or even slightly increase following drug treatment (**Supplementary** Fig.11). Overall, these findings suggest that Filgotinib mitigates the effect of limited resources, offering valuable insights for industrial applications.

### Improving viral vector transduction by Filgotinib supplementation

Gene and cell therapy has become a clinical reality with several market-approved *living drugs* for the treatment of monogenic diseases and hematological malignancies^38^. They are typically based on the genetic modification of the target cells with lentiviruses or AAV viruses. Lentiviruses are the primary choice to engineer eukaryotic cells *ex vivo* and are largely exploited for adoptive cellular therapy such as T cells engineered with chimeric antigen receptors (CARs) or cloned T-cell receptor (TCRs)^39^. However, transduction efficiency is hampered when lentiviruses include big genetic payloads, a limitation for complex regulatory network integration. Conversely, AAV derived vectors are used for *in vivo* therapy, can be engineered with tissue-specific tropism^40^ and ensure long-lasting transgene expression, making them suitable as delivery tool for therapeutic genes in several genetic disorders. However, a major limitation is that depending on the serotype, transduction of cells *ex vivo* may be inefficient, making circuits prototyping prior *in vivo* experiments largely ineffective.

We investigated whether Filgotinib would improve lentivirus and AAV transduction *in vitro,* since it demonstrated the most consistent effect across the different experimental settings. HEK293T cells were transduced with a lentivirus carrying a Ubc-EGFP and added the small molecule after 4h (**Fig.5a**). 72h post-Filgotinib treatment we observed ∼30% increase of EGFP-transduced cells as compared to untreated samples **(Fig. 5b-c)**.

**Figure 5.**
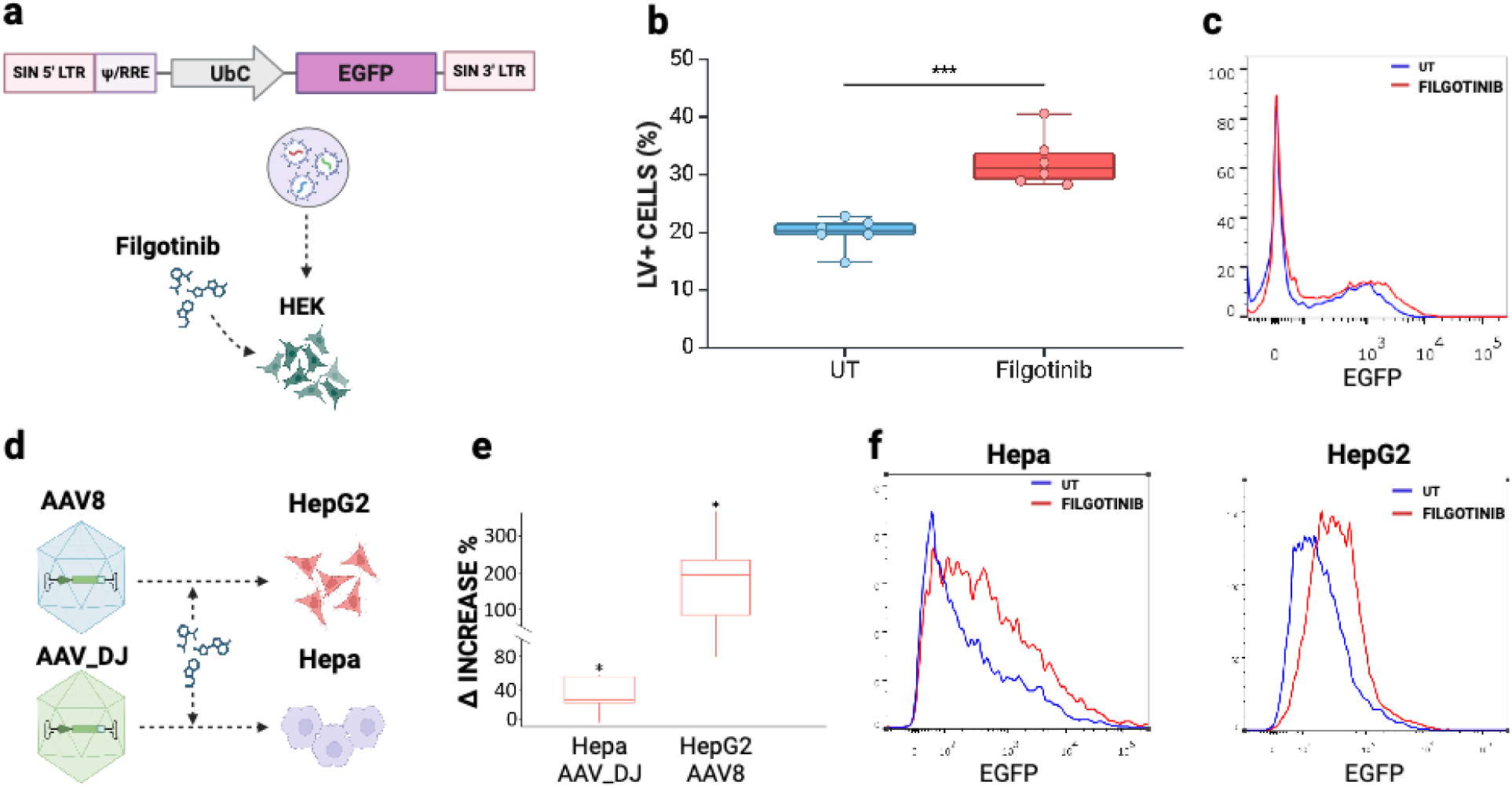
Enhancement of viral transduction by Filgotinib treatment. (**a**) Schematics of Filgotinib treatment during lentiviral transduction to improve efficiency in HEK293T cells. (**b**) Percentage of transduced cells in untreated and Filgotinib samples. (**c**) Exemplative histogram of lentiviral transduced cells w/wo Filfotinib. (**d**) Schematics of AAV transduction followed by Filgotinib treatment in hepatocyte-like cell lines Hepa and HepG2. (**e**) Percentage increase of AAV-positive cells after transduction and Filgotinib supplementation (**f**) Exemplative histogram displaying the differences in AAV expression w/wo Filgotinib. Paired T-test for statistical analysis (*p<0.05,**p<0.01,***p<0.001); (**b**) N=3 biological replicates. (**e-f**) N=4 biological replicates. *Created with BioRender.com*.

Next, to investigate the effect of Filgotinib on AAV transduction, we use hepatotropic AAVs, namely AAV8 and AAV_DJ, and we transduced HepG2 and Hepa1-6 (Hepa) hepatocytes. AAV8 is a natural AAV, while AAV_DJ is a type 2/type 8/type 9 chimera, distinguished from its closest natural relative (AAV-2) by 60 amino acids in the capsid^41^. In specific, HepG2 were transduced with AAV8 serotype expressing EGFP fluorescence, whereas Hepa cells deriving from murine hepatoma and less prone to AVV8^42,43^,were transduced with AAV_DJ that had already demonstrated to outperform standard AAV serotypes for in vitro transduction^41^ (**Fig.5d**). HepG2 treated with Filgotinib upon AAV8 transduction exhibited 100% increase of EGFP+ cells, whereas Hepa cells transduced with AAV_DJ exhibited a 40% increase of transduced cells (**Fig.5e-f**).

Last, we investigated the effect of Filgotinib supplementation on a population of HepG2 cells transduced by AAV_DJ (transduction efficiency close to 100%) and observed an increase of EGFP fluorescence of transduced cells as compared to the untreated cells (**Supplementary** Fig. 12). Collectively these data indicate that Filgotinib can be effectively used in a variety of applications for genetic design screenings and prototyping.

## Discussion

This study lays the groundwork of deploying computational algorithms for the identification and selection of small molecules that can reduce the impact of cellular burden and enhance exogenous gene expression, advancing industrial biomanufacturing and gene therapy vector prototyping. Cellular burden is a pervasive issue that affects the engineering of multiple organisms ranging from bacteria^44,45^ to mammalian cells^12,13^. Over the past few years, various strategies including genetic modules and alternative regulatory architectures have been proposed to reduce competition for the intracellular resources and to increase the cell operational capabilities^12–14^. By combining computational tools and experimental designs, we here provide a novel approach to increase cell productivity, which neglects both the modular composition of gene expression cassettes, and the network architecture. Small molecules can regulate several processes in the cells, including the production efficiency of exogenous DNA and RNA material. Through DECCODE algorithm^23^, we have identified compounds that upregulate protein expression with no prior information about the mode of action, avoiding biases on the bottlenecks of intracellular resources. Of the top sixteen drugs selected by DECCODE, Ruxolitinib, TWS119, Filgotinib and TIE2 inhibitor were exhibiting an enhancing effect on protein production and Filgotinib had consistent responses in a variety of different settings. Interestingly, we also observe that the drug treatment can support the optimization of gene expression of previously characterized genetic regulatory elements (e.g. HGHpolyA) (**Supplementary** Fig.2), opening further horizons of synergistic combinations to tune gene expression in a desired manner, facilitating applications across a broader spectrum of cell types and genetic constructs. Importantly, the impact of the drugs, similarly to the intracellular resource competition, is cell context dependent. For example, CHO cells that respond to drugs such as rapamycin ^46,47^, do not increase gene expression with the tested drugs. This may be due to different origin of the cell line or different metabolism than the human-derived cell lines. The highest increase of transgene expression was achieved with Filgotinib and Ruxolitinib, the first emerging as lead candidate across several settings. The effect of Filgotinib on viral preparation and transduction can significantly improve the scalability and efficiency of vector screening and production, holding promising applications in biomanufacturing and gene therapy. For instance, the viral production costs can be drastically reduced by increasing the titer with the compound supplementation. Filgotinib and Ruxolitinib target the JAK-STAT pathway. Filgotinib in specific inhibits JAK1, preventing the activation of STATs, ultimately leading to a reduction in proinflammatory cytokine signaling ^46–47^. Although understanding the mechanism by which protein production increases is not a goal of this work, we were curious to understand whether the augmented cells operational capacity might be a consequence of this inhibition. We silenced JAK1 expression in H1299-EGFP cell line using RNA interference (RNAi) with a JAK1-specific siRNA and monitored EGFP expression, mimicking the condition of **Fig.4b**. While we successfully silenced the JAK1 gene, the results did not replicate the effects observed with Filgotinib treatment (**Supplementary** Fig. 13 **left**). This suggests that the increase in productivity may not be directly linked to JAK1 inhibition alone. However, it was interesting to observe that also the expression of the JAK1 transcripts increased by Filgotinib supplementation, supporting the insights of a global enhancement of genes’ expression (**Supplementary** Fig. 13 **right**). This effect was similar to what observed with miRNA-based iFFL^15^. We carried out a DSEA analysis on transcriptomic data of the four validated drugs, to examine whether enriched pathways may relate to those that resulted dysregulated by the transcriptomics of the burdened cells (**Fig. 2c**). We found an enrichment of pathways that relate to RNA stability and maturation, ribosome assembly, and tRNA metabolism, that suggest a general impact on the post-transcriptional processes and translation (**Supplementary** Fig.14). Others are involved in the metabolism suggesting an effect also on energy production and cellular activity (**Supplementary** Fig.14). In perspective, understanding the exact mechanism of action of the drugs on the cellular machine will allow to identify specific genetic targets for the engineering of cells with enhanced productive capabilities.

In summary, our study demonstrates the power of combining computational algorithms with synthetic devices to increase cellular productivity without needing detailed prior knowledge of drug mechanisms or resource competition bottlenecks. This approach has significant potential for *in vitro* systems and could pave the way for more efficient biomanufacturing and rapid prototyping of genetic circuits for therapeutic applications.

## Materials and Methods

### Cell culture

HEK293T, U2OS, MDA and Hepa cells used in this study were cultured in Dulbecco’s modified Eagle medium (DMEM, Gibco); H1299 were cultured in Roswell Park Memorial Institute medium (RPMI, Gibco); CHO-K1 were cultured in α-MEM (Sigma Aldrich, M4526); HepG2 were maintained in MEM-E. All media were supplemented with 10% FBS (Atlanta BIO), 1% penicillin/streptomycin/L-Glutamine (Sigma-Aldrich) and 1% non-essential amino acids (HyClone). The cells were maintained at 37 °C and 5% CO2.

### Stable cell lines engineering

H1299-EGFP, HEK-EGFP, MDA-MB231-EGFP and U2OS-EGFP were generated by transduction with a lentivirus carrying an EGFP gene under the UbC promoter. HEK-mCherry were obtained by transducing cells with a lentivirus carrying a shEF1α-mCherry construct. In all the cases 8*10^5 cells were plated in a 6 multiwell, and the virus was added in a ratio 1:3 with the culture medium 6h after the plating. Cells were analyzed one week after transduction to assess the fluorescence and then maintained and amplified as described previously.

CHO-TET-OFF cells (Clontech #630904) transduced with a lentivirus containing CMV-TET promoter that drives the expression of tTA and d2EYFP described in Siciliano et al. paper^27^ were a kind gift from Diego di Bernardo (Telethon Institute of Genetics and Medicine-TIGEM, Naples, Italy).

### Transfection

Transfections were carried out in 24-well plate for flow cytometry analysis. H1299 cells were transfected with Lipofectamine® 3000 according to manufacturer’s instructions using 300ng of total DNA in 24-well plates. CHO and HEK293T were transfected with PEI transfection reagent with 300 ng of total DNA in 24-well plates. 70000 to 80000 cells per well were plated approximately 24h before transfection. At the moment of transfection, cells were put in starvation conditions for the first 24h. DNA was diluted in Opti-MEM reduced serum media (Gibco), before being mixed and incubated for 25 minutes prior to addition to the cells. All drugs were added 4h after transfection.

Modified-RNAs (modRNA) were transfected using Lipofectamine® 3000 protocol under modified conditions in which no P3000 reagent was used. RNA was first diluted in Opti-MEM reduced serum media (Gibco) and it was then mixed with diluted Lipofectamine reagent as per manufacturer’s instructions. The mix was incubated for 7 minutes prior to addition to the cells. The fast-forward protocol was used by seeding 120000 cells per well in 24-wells plates at the moment of transfection.

### modRNA production

Modified-RNAs were produced by in vitro transcription (IVT) performed using MegaScript T7 kit (Life Technologies). In the IVT reactions, modified conditions were used: GTPs were replaced by GTPs mixed with Anti Reverse Cap Analog (New England BioLabs) at the ratio of 1 to 4; while CTPs and UTPs were replaced by 5-methylcytosine-triphosphate and pseudouridine-triphosphate (TriLink BioTechnologies), respectively. Transcripts were then treated with Turbo DNase included in MegaScript T7 kit (Life Technologies) for 30 min at 37 °C and purified using MEGAclear™ Transcription Clean-Up Kit (Life Technologies). Lastly, resulting modified mRNAs (modRNAs) were incubated with Antarctic Phosphatase (New England BioLabs) for 30 min at 37 °C and purified again.

### Drug preparation

All small molecules are commercially available and purchased by Selleck except for TWS119 that is from VWR. All compounds are dissolved in DMSO. Ruxolitinib and Filgotinib are used 10uM^27,29^. TWS119 is used 2uM^28^ and TIE2 inhibitor is used 5uM^30^.

### Lentivirus production

Lentivirus productions were carried out in 6-well plate in Hek293T cells. HeK293T were transfected with second-generation lentiviral plasmids with transfer vector and packaging plasmids in a ratio of 2:1:1 using the transfection method described above. Drugs were added 4h after transfection. Media was changed 24h after transfection with complete media and no drugs was added afterwards. The virus was collected 72h after transfection and concentrated with Lenti Concentrator (OriGene technologies) following manufacturer instructions. Cell lines were transduced with same MOI.

### Lentivirus and AAV transduction

Lentivirus and AAV transduction were carried out in 24-well plates. Lentivirus was added to cells at the dilution of 1:2000. The AAV are used at an MOI of 1×10e^5^ gc/cell. The virus is added to incomplete media, incubated for 2h and 30 min, and then added to plated cells. Drugs were added 4h after AAV infection.

### Lentivirus titer

Lentivirus titrations were carried out in 24-well plates. Lentivirus was added to cells at the dilution of 1:50; 1:100; 1:200; 1:500.

After 72h the EGFP fluorescence was measured by flow cytometry and based on the samples with a percentage of fluorescence of 20-40% titer was calculated with this formula:

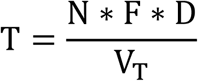

where T = Titer, TU/mL; N = Number of cells transduced; F = Fraction of cells with fluorescence; D = Dilution Factor; V_T_ = Transduction Volume, ml^32^.

### AAV Production

AAVs were provided by Nicola Brunetti Pierri (Telethon Institute of Genetics and Medicine-TIGEM, Naples, Italy) The pAAV2.TBG.GFP^50^ was used for production of both AAV_Dj and AAV_8 serotype vectors. AAV vectors were produced by Innovavector s.r.l., Italy, by triple transfection in HEK293T cells; next were purified by CsCl2 ultracentrifugation and titered (in genome copies/milliliter) by real-time PCR and dot blot analysis.

### Flow cytometry and data analysis

All cells were analyzed with a BD CELESTA™ cell analyzer (BD Biosciences). For each sample >10000 singlet events were collected. Cells transfected in 24-well plate were washed with DPBS, detached with 50 μl of Trypsin-EDTA (0.25%) and resuspended in 300 μl of FACS medium. Fluorescence intensity in arbitrary units (au) was used as a measure of protein expression. For each experiment a compensation matrix was created using unstained (wild type cells), and single-color controls (mKate only, EGFP only). Population of live cells and single cells were selected according to FCS/SSC parameters. Data analysis was performed with FlowJo.

### ELISA assay

ELISAs are performed with the Human Secondary Lymphoid-tissue Chemokine (CCL21) (SLC) Uncoated ELISA Kit (ThermoFisher Scientifics). Supernatant were harvested from 24 well plates with transfected H1299 or HEK293T cells 48 hours after transfection and CCL21 was quantified according to manufacturer instructions.

### Luminescence assay

Luminescence assays were performed on transfected HEK293T cells 48h after transfection with split luciferase to detect luciferase activity. The luciferase activity was performed by Steady-Glo Luciferase Assay System (Promega). A volume of SteadyGlo reagent was added to 100ul of cultured cells in a white 96 multiwell plate, following the manufacturer instructions 5 minutes after the addiction of the reagent we measured the luminescence on Glomax multimode microplate reader and normalized by the number of cells counted prior to the assay.

### RNA-seq samples preparation

For RNA-seq analysis Qiagen RNeasy mini plus Kit (Qiagen) was used for RNA isolation. 10 µL of RNA at the exact concentration of 50 ng*µL −1 were provided for library preparation to the NGS facility of the Telethon Institute for Genetis and Medicine-TIGEM (Naples, Italy). Libraries were prepared using QuantSeq 3’ mRNA sequencing for RNA quantification kit (Lexogen). Samples were processed with NovaSeq 6000 System (Illumina).

### RNA-Seq Analysis

Sequence reads were trimmed using bbduk software (https://www.lexogen.com/quantseqdata-analysis/) to remove adapter sequences, poly-A tails and low-quality end bases. Alignment was performed with STAR^51^ on the Homo sapiens reference (hg38) provided by UCSC Genome Browser. The expression level of genes was determined with htseq-count^52^ using the Gencode/Ensembl gene model. Differential expression analysis was performed using DESeq2^53^, a statistical package based on a model using the negative binomial distribution. Only genes with FDR 0.1 were considered differentially expressed for each comparison and were used to determine whether known biological functions or processes are enriched by clusterProfiler package^54^.

### DECCODE analysis

The Gep2Pep R/Bioconductor package^55^ was employed to obtain the pathway-based expression profiles of the using 14,645 gene sets from 16 different gene set collections included in the MsigDB v6.1^56^. This version was chosen to match the precomputed pathway-based version of the LINCS drug-induced profiles dataset that is publicly available^55^. As previously described^22^, the gene set database retaining most of the gene-based information (i.e., “Gene Ontology - Biological Process”) was selected for further analysis. Finally, the DECCODE method was applied to prioritize pathway based LINCS^24^ profiles based on their similarity to the target signature. The similarity scores were obtained as the reciprocal of the Manhattan distance between each profile pair after ranking pathways by their Enrichment Scores.

### Statistics and reproducibility

Each experiment was repeated independently at least three times with similar results. To compare multiple drugs, statistical analysis was performed using a randomized block design ANOVA with two additive factors: treatment and experiment. The treatment factor, which was the factor of interest, consisted of the different drugs used to treat the cells. The experiment factor, known as the block, was used to control a known source of variability and its levels were constituted by the different experiments. This was followed by a Dunnett post hoc test to identify significant differences among groups. For comparisons between two conditions, a two-tailed paired t-test was used. Both ANOVA and t-test were performed on the log-transformed data. In the figures, we represent the percentage increase, or the fold change calculated on the log-transformed data, comparing the drug treated with the untreated sample. To calculate the percentage increase, we used the formula:

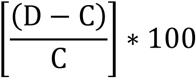

To calculate the fold change, we used the formula:

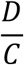

In both cases D is the Drug treated sample and C is the control (untreated sample).

The threshold for significance was set to 0.05 for the global ANOVA and t-test p-values, as well as for the adjusted p-values in the post hoc analysis. Prior to any testing, we assessed the normality of the data using the Shapiro-Wilk test and checked for homogeneity of variance using the Levene’s test.

## Data availability

All relevant data are included as Source Data and/or are available from the corresponding author on reasonable request. Plasmids used in this study are available from the corresponding author on reasonable request.

## Code availability

The authors are confident that the conclusions do not strongly depend on the particular choice of analysis software. Nevertheless, the code used for automated analysis and fitting is available on reasonable requests from the corresponding authors.

## ACKNOWLEDGEMENTS

We thank Daniela Perna for technical support. We also thank the genomic facility of IIT for the RNA sequencing support. Author contributions: V.S. conceived the project, V.S. and M.P. designed experiments. M.P. performed the experiments. F.C. performed the transfection for RNA-sequencing. F.CA. designed and produced the modified RNA. I.B. and N.B.P. produced the AAV vectors. A.R. analysed the RNA-seq data and provided bioinformatics support. A.C., M.P., and V.S. analysed the data. A.C. performed statistical analysis. F.T. performed experiments in integrated cell lines. F.N. performed the computational work. V.S. supervised the experimental work and secured funding. M.P., F.N,. A.R., V.S. wrote the manuscript. F.C., A.C. edited the manuscript.

