## Supplementary Information for "Computational identification of small molecules for increased gene expression by synthetic circuits in mammalian cells"

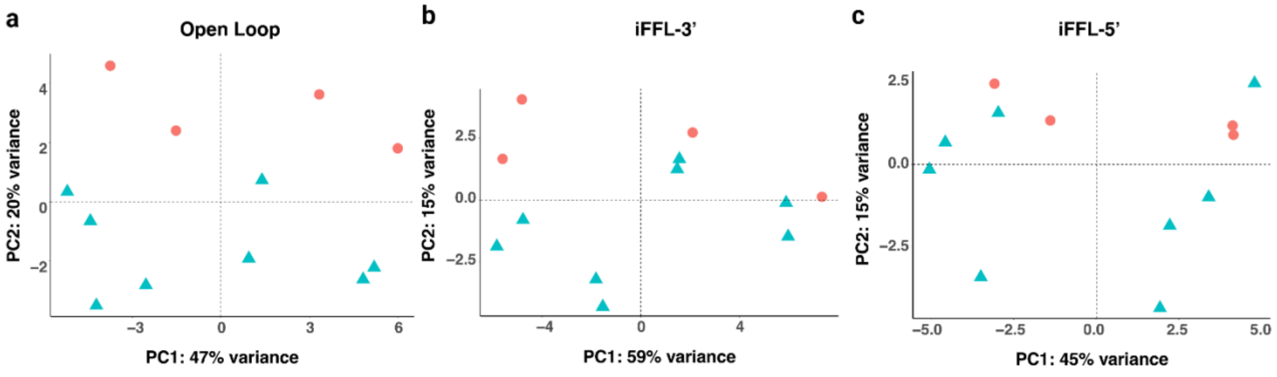


**Supplementary Figure 1 PCA of RNA-Sequencing data.** (**a**) PCA of the Open Loop (OL) topology showing that the non-transfected population (NTr-red dots) is spatially separated from the transfected population (Tr-blue dots), while in the iFFL-3’ (**b**) and iFFL-5’ (**c**) this difference is progressively reduced. Data were collected 48h post-transfection. N = 4-8 biological replicates.

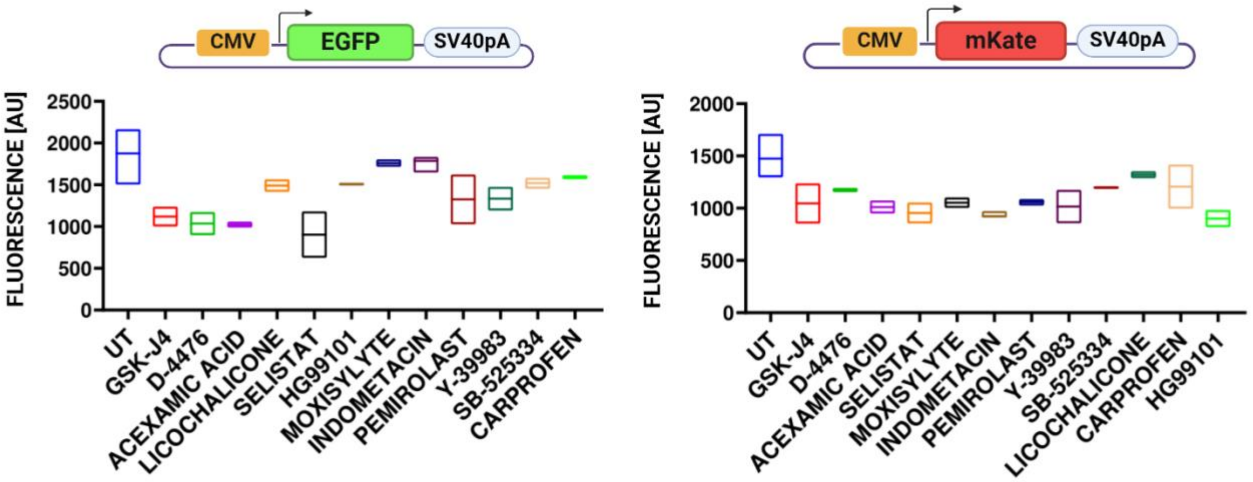


**Supplementary Figure 2 Validation of compounds identified by DECCODE in H1299 cell line.** Geometric mean fluorescence of samples treated with the top sixteen drugs identified by the DECCODE, indicating that the drugs did not show increase of EGFP or mKate fluorescence compared to untreated samples. Source data are provided as a Source Data file. N=2 biological replicates for each condition.

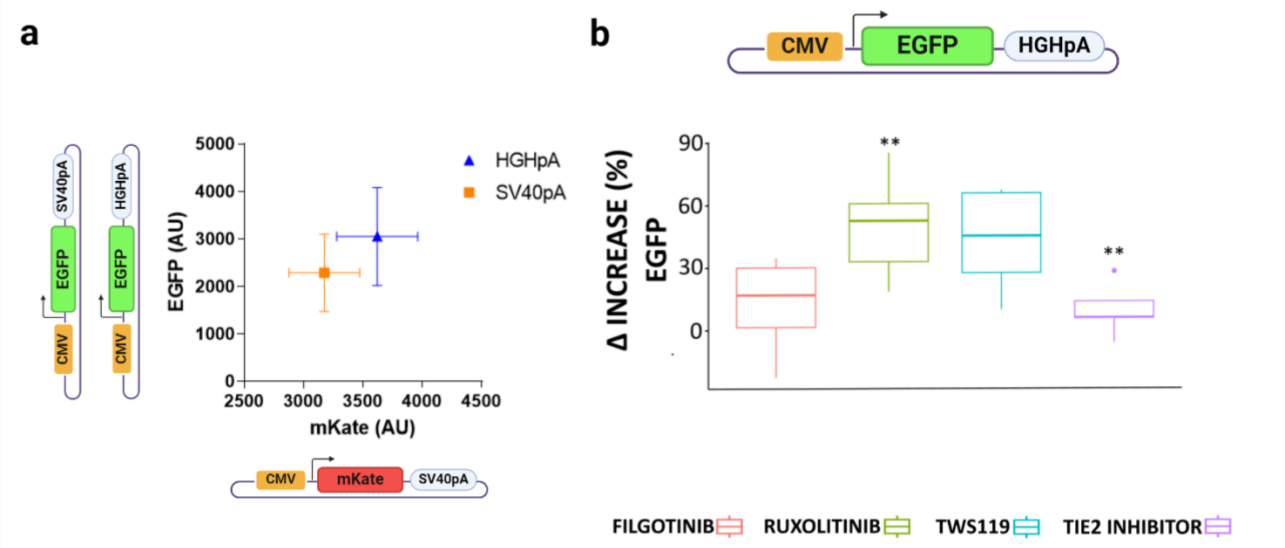


**Supplementary Figure 3 Response to drugs of different genetic modules composition.** (**a**) impact of the different genetic modules on protein expression in H1299 cells. As previously demonstrated for HEK293T and CHOK1[1], the plasmids with HGHpolyA is confirmed to be the best in terms of expression of both plasmids at the same time. Error bars show standard error of n=5 biological replicates (**b**) Percentage increase in fluorescence intensity (delta increase) for EGFP-HGHpA in response to four different drugs: Filgotinib, Ruxolitinib, TWS119, and a TIE2 inhibitor. ANOVA test for statistical analysis (*p<0.05,**p<0.01,***p<0.001); N=5 biological replicates.

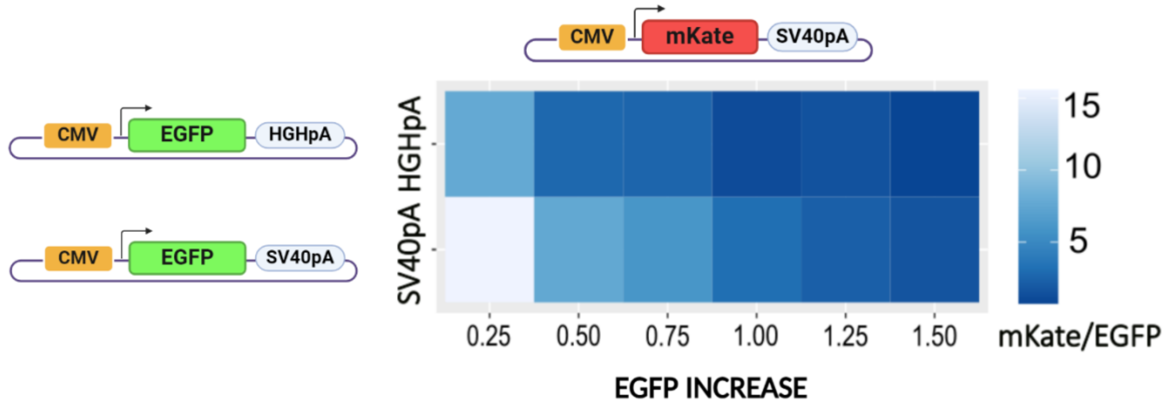


**Supplementary Figure 4 Response of different genetic modules combination to gene perturbation.** Increasing EGFP-HGHpA or EGFP-SV40pA encoding plasmid was provided to induce burden on mKate provided at the same amount similar to what was done in Frei et al work[2]. The x-axis represents the molar ratio of EGFP:mKate. The color intensity on the heatmap indicates the mKate/EGFP ratio across the different molar ratio is consistently lower in the combination EGFP-HGHpA/mKate-SV40pA.

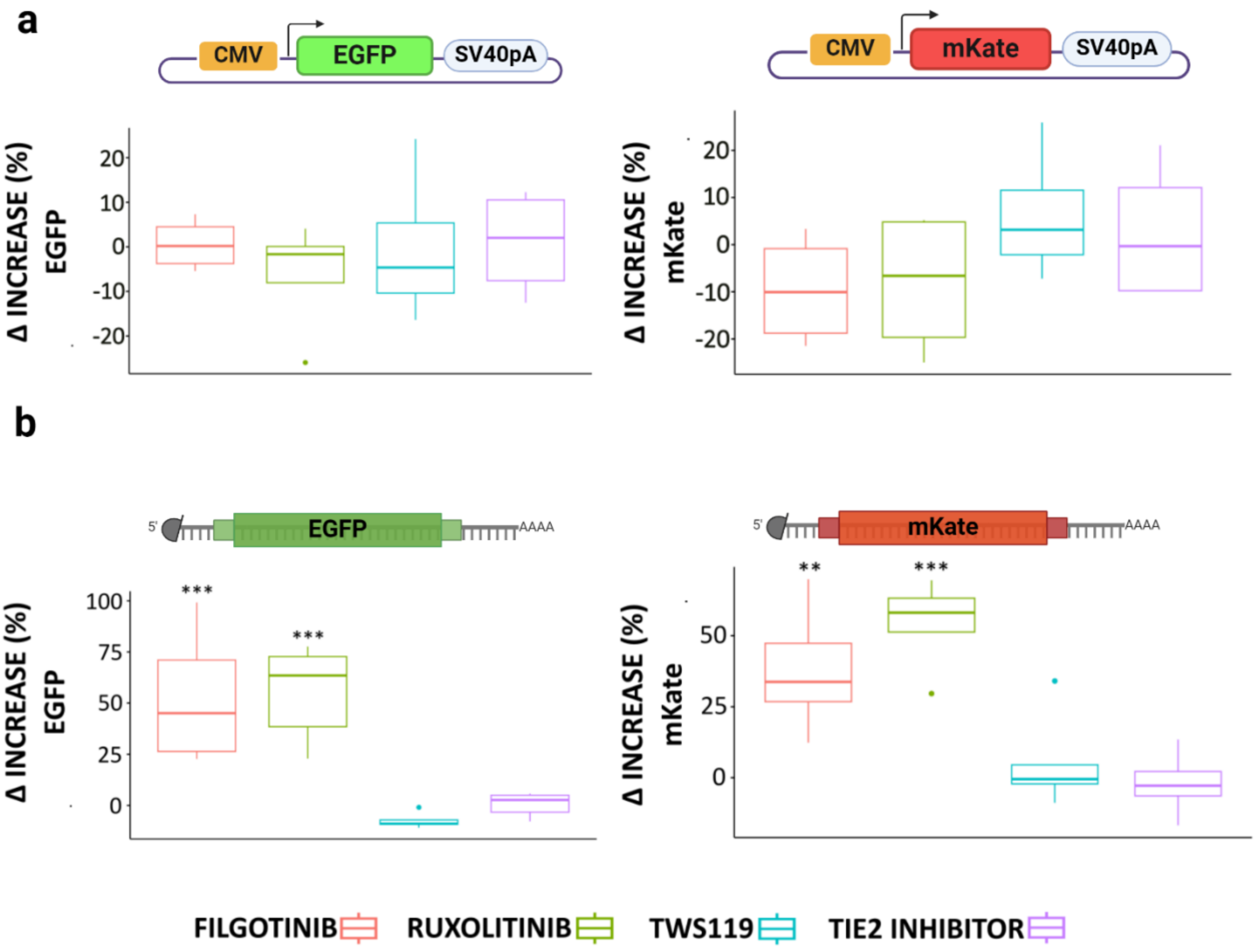


**Supplementary Figure 5 CHO-K1 cell line response to drugs.** (**a**) Response to drug treatment on DNA-encoded EGFP and mKate. The boxplots representing fluorescence increase as compared to untreated samples show no significant difference in fluorescence intensity. (**b**) Drugs treatment on RNA-encoded EGFP and mKate results in a 30 to 60% increase of fluorescent protein expression. (**a**) N=4 biological replicates. (**b**) N=5 biological replicates. ANOVA test for statistical analysis (*p<0.05,**p<0.01,***p<0.001).

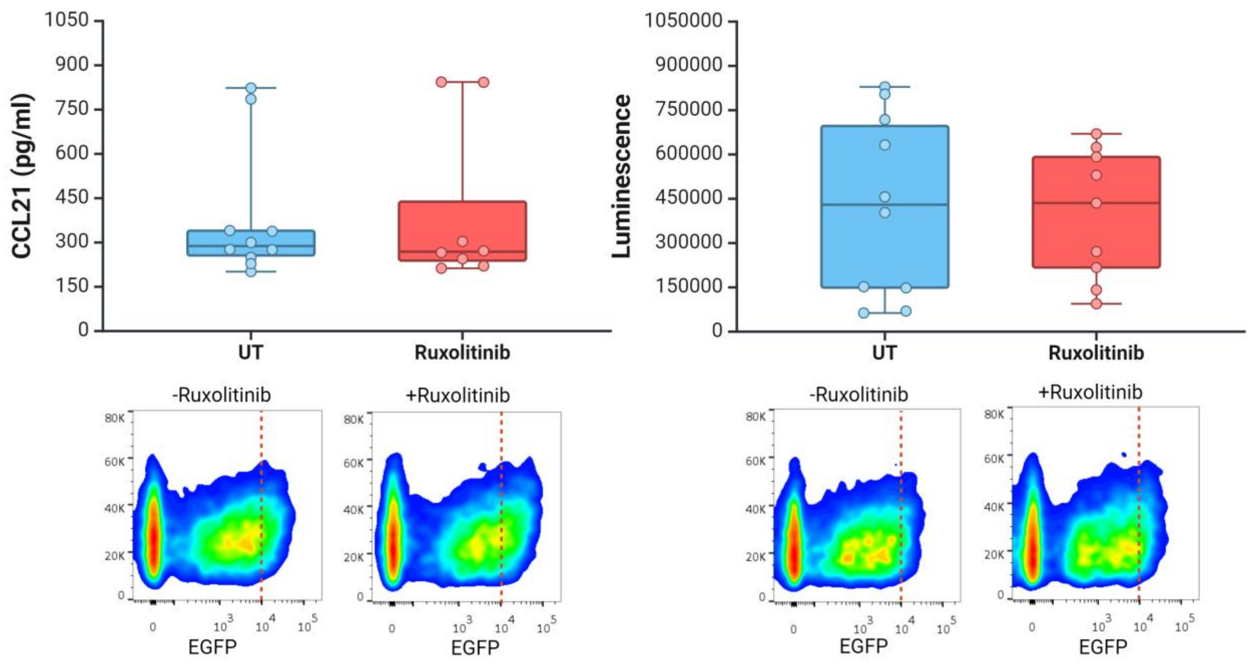


**Supplementary Figure 6 CCL21 and Luciferase expression after Ruxolitinib treatment.** Ruxolitib treatment does not increase the chemokine CCL21 and the Luciferase expression. The boxplots show the concentration per million of cells in CCL21 expression and luminescence signal following drug treatment. The density plots display EGFP fluorescence (co-expressed with CCL21 or with split-Luc) with and without Ruxolitinib treatment. Boxplots of n=5 biological replicate. *Created with BioRender.com.*

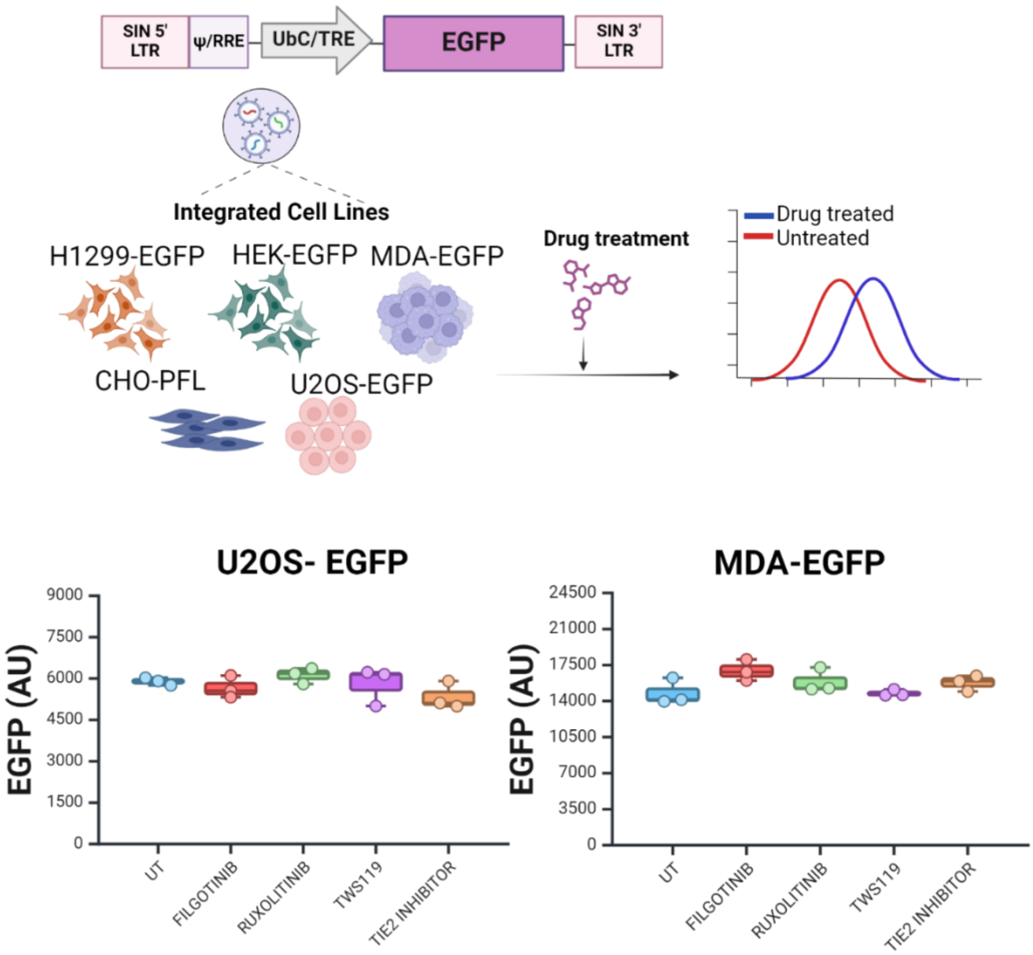


**Supplementary Figure 7 Response to drugs of stably integrated genes in different cell lines. Top**: experimental pipeline for stably cell engineering. EGFP was integrated by lentiviral transduction, followed by treatment with the small molecules and fluorescence intensity assessment. **Bottom**: geometric mean (AU, arbitrary units) of EGFP in untreated samples (UT) and samples treated with the four different drugs. The results indicate that neither U2OS-EGFP nor MDA-EGFP cell lines show significant changes in fluorescence intensity. Boxplots represent geometric mean of N=3 biological replicates. *Created with BioRender.com.*

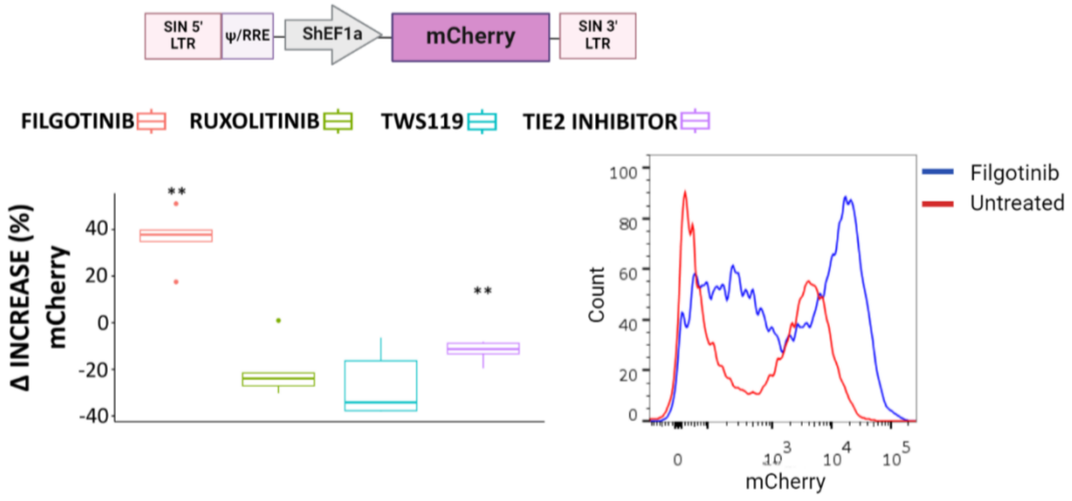


**Supplementary Figure 8 HEK293T-mCherry cell line response to drug. Top**: lentiviral vector with a shEF1a promoter driving the mCherry reporter. **Bottom**: delta increase percentage in fluorescence intensity of mCherry in response to the four different drugs compared to untreated samples. The data show a 40% increase in fluorescence intensity in HEK293T cells upon Filgotinib supplementation. On the right, histogram representing HEK293T-mCherry with and without Filgotinib treatment. This suggests enhanced protein production in response to Filgotinib treatment across different promoter configurations. N=5 biological replicate. ANOVA test for statistical analysis (*p<0.05,**p<0.01,***p<0.001).

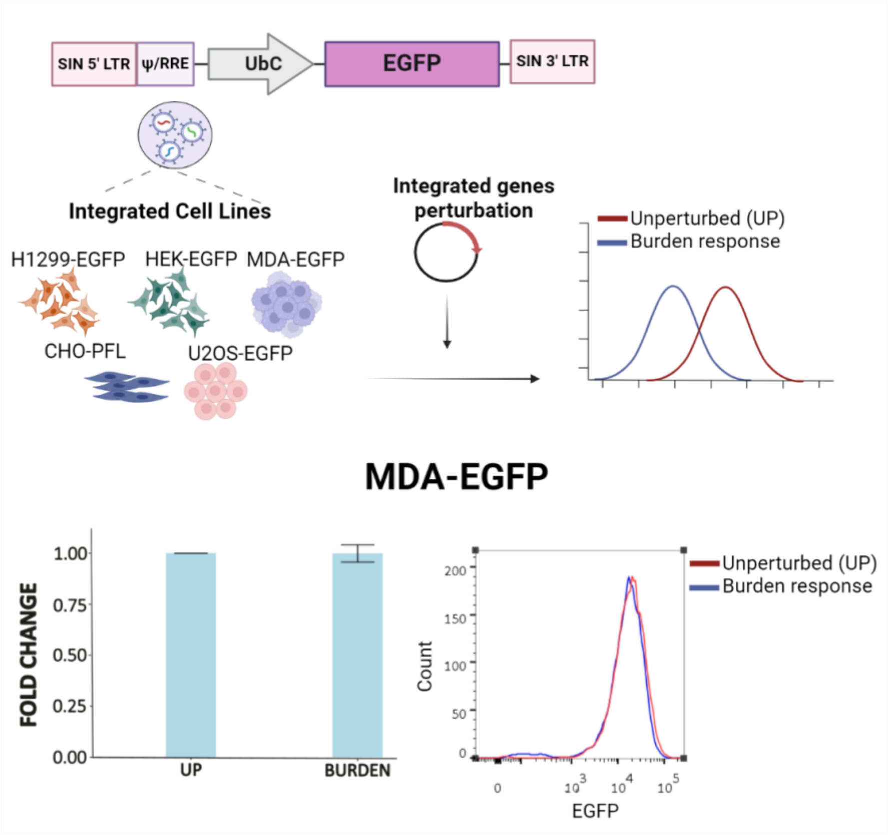


**Supplementary Figure 9 Integrated payloads perturbation by transient transfection in MDA-EGFP cell line. Top**: experimental pipeline used to assess the response of integrated payloads to transient transfection perturbations. **Bottom**: bar plots represent the fold change expression of EGFP upon CMV-mKate transient expression, showing that MDA-EGFP cells are resilient to burden. On the right, histogram representing MDA-EGFP with and without perturbation. N=4 biological replicates. Error bars represent standard error, SE. *Created with BioRender.com.*

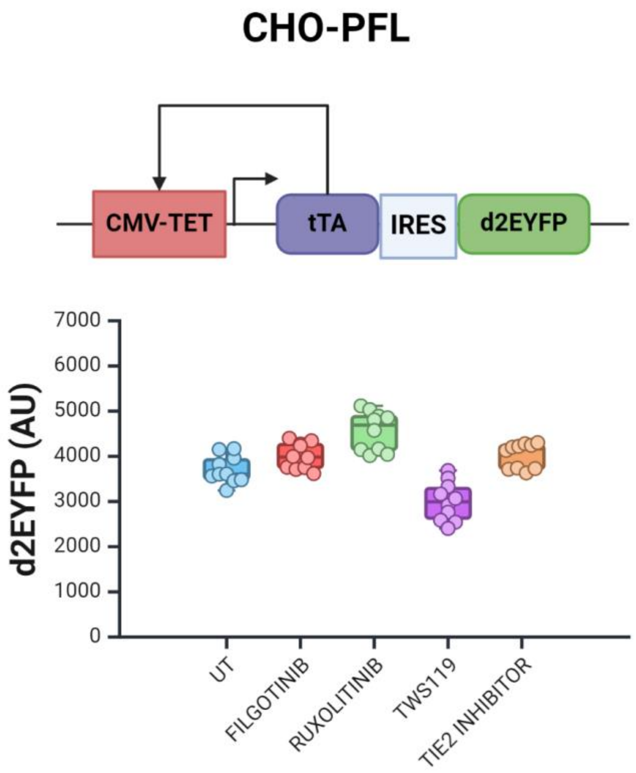


**Supplementary Figure 10 CHO-PFL cell line response to drug. Top**: schematics of the positive feedback loop-PFL circuit integrated in CHO-TET-OFF developed in Siciliano V et al 2013[3]. **Bottom**: box plots comparing the geometric mean (AU, arbitrary units) of d2EYFP in untreated samples (UT) and samples treated with four different drugs. The results indicate that CHO PFL cells do not show significant changes in fluorescence intensity in response to any of the tested drugs. N=5 biological replicates. *Created with BioRender.com*

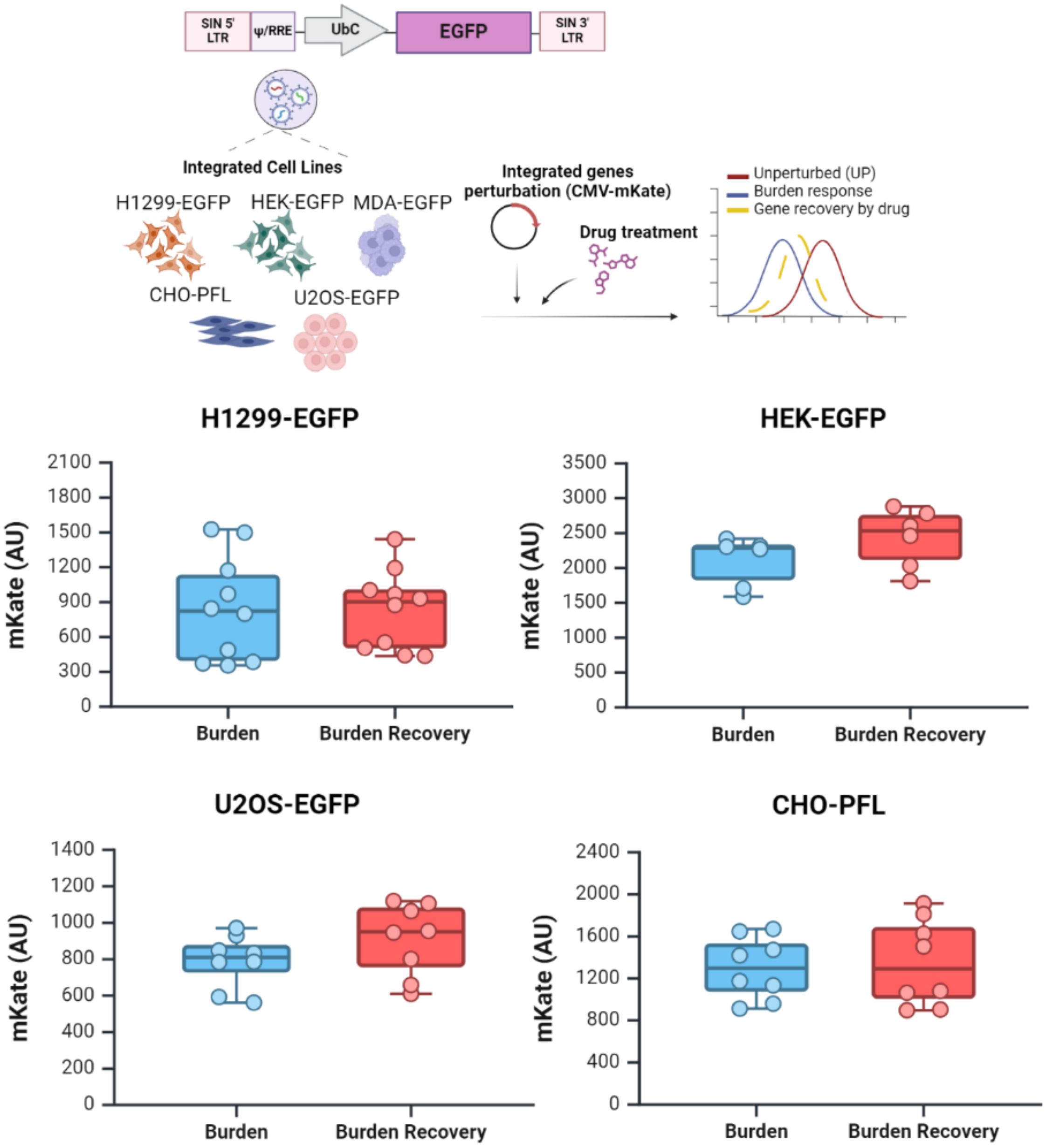


**Supplementary Figure 11 mKate exogenous expression in integrated cell lines. Top**: experimental pipeline used to assess the response of integrated payloads to transient transfection perturbations and drug treatment. **Bottom**: geometric mean of mKate used to impose burden in untreated samples (Burden) or in Filgotinib treated samples (Burden recovery) in the target cell lines. N=4 biological replicates. *Created with BioRender.com.*

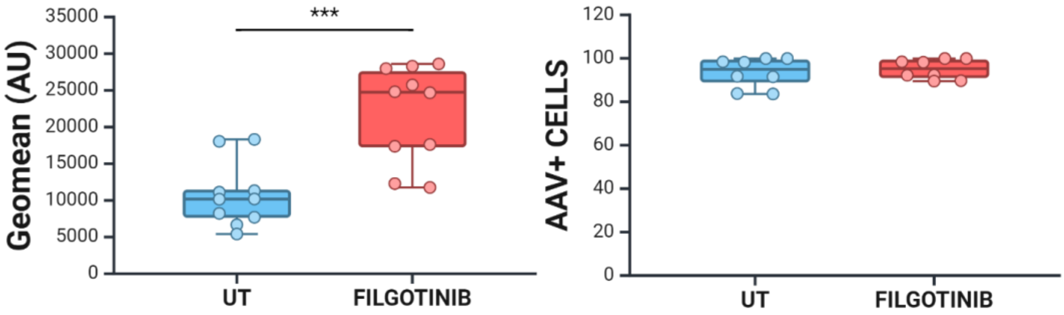


**Supplementary Figure 12 HepG2 transduction with AAV_DJ serotype. Left**: Filgotinib treatment resulted in a significantly higher geometric mean fluorescence compared to the untreated group. **Right**: percentage of AAV_DJ transduced HepG2 already near the 100% did not improve by small drug supplementation. N=5 biological replicates. Unpaired t-test for statistical analysis (*p<0.05,**p<0.01,***p<0.001). *Created with BioRender.com.*

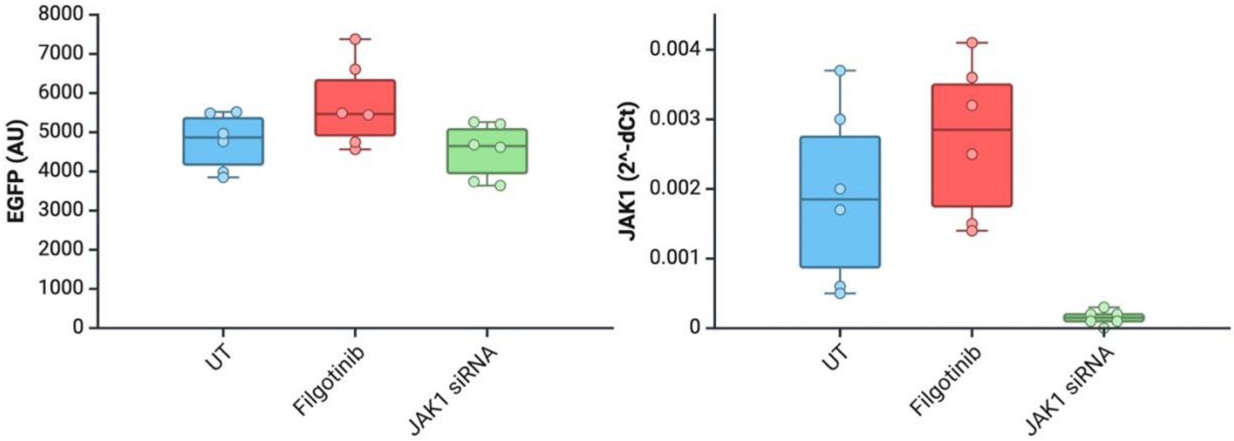


**Supplementary Figure 13 Effect of JAK1 siRNA on EGFP Expression in H1299-EGFP. Left**: Box plots representing the geometric mean of EGFP showing the expected increase in EGFP expression in Filgotinib-treated sample, while the treatment with JAK1 siRNA do not show any results. **Right**: mRNA expression ($2^{\text{-dCt}}$, right panel) in untreated (UT), Filgotinib-treated, and JAK1 siRNA-treated samples, displaying JAK1 mRNA expression quantified by qPCR. N=3 biological replicates. *Created with BioRender.com.*

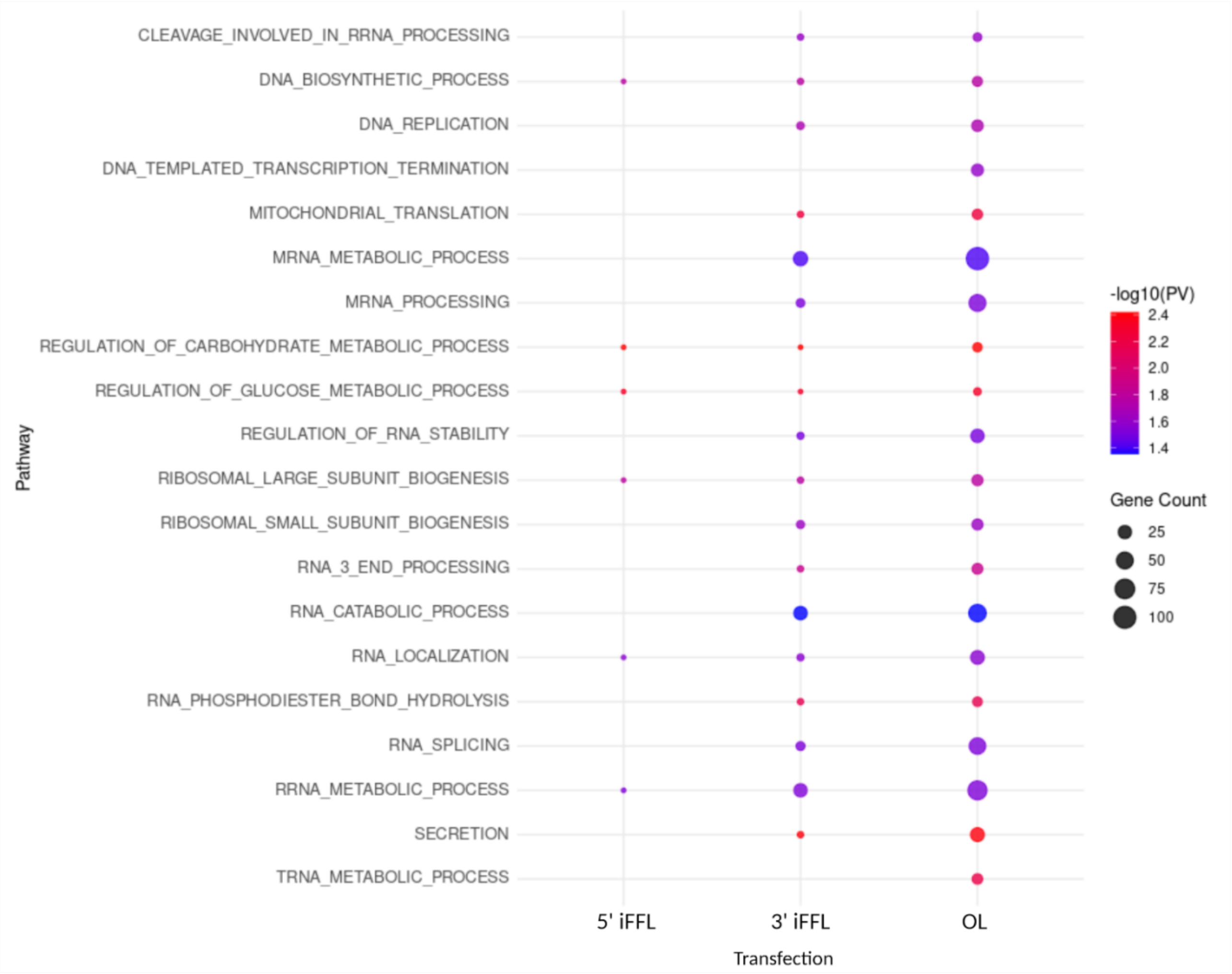


**Supplementary Figure 14. Drug-set Enrichment Analysis (DSEA) and Pathways over-representation analysis (ORA).** Bubble plot showing the common biological pathways between DSEA of the four validated drugs and ORA across the three different transfections (iFFL3, iFFL5, and OL). The pathways are listed on the y-axis, while the transfections are categorized on the x-axis. The size of each bubble indicates the gene count associated with each pathway, with larger bubbles representing a higher gene count. The color intensity of the bubbles represents the statistical significance level of DSEA, with colors ranging from blue (less significant) to red (more significant) based on the –log10 (PV). Notably, pathways with high gene count and significant p-values include mRNA metabolic process, ribosomal large subunit biogenesis, and tRNA metabolic process among others.

| Sample | FDR<=0.1 | LogFC >=0.5 | LogFC <=-0.5 |
|---|---|---|---|
| OL | 1088 | 155 | 109 |
| iFFL-5’ | 18 | 0 | 8 |
| iFFL-3’ | 111 | 6 | 1 |

**Table 1.** Number of DEGs in each transfection condition according to the RNA-seq analysis.

| Sample | GOBP | GOCC | GOMF |
|---|---|---|---|
| OL | 158 | 80 | 38 |
| iFFL-5’ | 4 | 0 | 0 |
| iFFL-3’ | 10 | 16 | 5 |

**Table 2.** Number of enriched pathways for each condition. GOBP = Gene Ontology Biological Process. GOCC = Gene Ontology Cellular Component. GOMF = Gene Ontology Molecular Factor.
